# A complete high quality nanopore-only assembly of an XDR *Mycobacterium tuberculosis* Beijing lineage strain identifies novel variation in repetitive PE/PPE gene regions

**DOI:** 10.1101/256719

**Authors:** Arnold Bainomugisa, Tania Duarte, Evelyn Lavu, Sushil Pandey, Chris Coulter, Ben J. Marais, Lachlan Coin

**Author notes:** **Corresponding Authors:** 1. Arnold Bainomugisa, Institute for Molecular Biosciences, The University of Queensland, St Lucia, Brisbane, Australia; 2. Assoc. Professor Lachlan Coin, Institute for Molecular Biosciences, The University of Queensland, St Lucia, Brisbane, Australia.

## Abstract

A better understanding of the genomic changes that facilitate the emergence and spread of drug resistant *M. tuberculosis* strains is required. Short-read sequencing methods have limited capacity to identify long, repetitive genomic regions and gene duplications. We sequenced an extensively drug resistant (XDR) Beijing sub-lineage 2.2.1.1 “epidemic strain” from the Western Province of Papua New Guinea using long-read sequencing (Oxford Nanopore MinION®). With up to 274 fold coverage from a single flow-cell, we assembled a 4404947bp circular genome containing 3670 coding sequences that include the highly repetitive PE/PPE genes. Comparison with Illumina reads indicated a base-level accuracy of 99.95%. Mutations known to confer drug resistance to first and second line drugs were identified and concurred with phenotypic resistance assays. We identified mutations in efflux pump genes (Rv0194), transporters (*secA1*, *glnQ*, *uspA*), cell wall biosynthesis genes (*pdk*, *mmpL*, *fadD*) and virulence genes (*mce*-gene family, *mycp1*) that may contribute to the drug resistance phenotype and successful transmission of this strain. Using the newly assembled genome as reference to map raw Illumina reads from representative *M. tuberculosis* lineages, we detect large insertions relative to the reference genome. We provide a fully annotated genome of a transmissible XDR *M. tuberculosis* strain from Papua New Guinea using Oxford Nanopore MinION sequencing and provide insight into genomic mechanisms of resistance and virulence.

**Data Summary:**

1. Sample Illumina and MinION sequencing reads generated and analyzed are available in NCBI under project accession number PRJNA386696 (https://www.ncbi.nlm.nih.gov/sra/?term=PRJNA386696)
2. The assembled complete genome and its annotations are available in NCBI under accession number CP022704.1 (https://www.ncbi.nlm.nih.gov/sra/?term=CP022704.1)

**Impact statement:** We recently characterized a Modern Beijing lineage strain responsible for the drug resistance outbreaks in the Western province, Papua New Guinea. With some of the genomic markers responsible for its drug resistance and transmissibility are known, there is need to elucidate all molecular mechanisms that account for the resistance phenotype, virulence and transmission. Whole genome sequencing using short reads has widely been utilized to study MTB genome but it does not generally capture long repetitive regions as variants in these regions are eliminated using analysis. Illumina instruments are known to have a GC bias so that regions with high GC or AT rich are under sampled and this effect is exacerbated in MTB, which has approximately 65% GC content. In this study, we utilized Oxford Nanopore Technologies (ONT) MinION sequencing to assemble a high-quality complete genome of an extensively drug resistant strain of a modern Beijing lineage. We were able to able to assemble all PE/PPE (proline-glutamate/proline-proline-glutamate) gene families that have high GC content and repetitive in nature. We show the genomic utility of ONT in offering a more comprehensive understanding of genetic mechanisms that contribute to resistance, virulence and transmission. This is important for settings up predictive analytics platforms and services to support diagnostics and treatment.

## Introduction

Globally, the tuberculosis (TB) incidence rate has shown a slow decline over the last two decades, although absolute case numbers continue to rise due to population growth, with an estimated 10.4 million new cases occurring in 2016 (1). TB control gains are threatened by the growing number of drug resistant strains recorded around the world (2). An estimated 490,000 incident cases of multi-drug resistant (MDR) TB, which is resistance to at least isoniazid and rifampicin occurred in 2016, accounting for 374,000 deaths among HIV-positive patients (1). The incidence of extensively drug resistant (XDR) strains, which are MDR strains with additional resistance to at least one fluoroquinolone and second line injectable, is also on the rise (1). Further multiplication of drug resistance in strains that are already highly drug resistant could lead to programmatically incurable TB, where construction of a curative regimen might be impossible with existing treatment options (3, 4). Management of drug resistant tuberculosis places a major financial burden on health systems, which may be overwhelmed in settings with high disease burdens (3).

In the absence of lateral gene transfer (5), drug resistance in *M. tuberculosis* arises mainly from chromosomal mutations that are selected by chemotherapeutic pressure, which drives drug resistance multiplication and the ongoing evolution of drug resistant strains (6-8). Successful transmission of drug resistant strains results in clonal expansion and potential epidemic spread (9-11). The acquisition of resistance-conferring mutations has potential for epidemic spread if these drug resistant strains are readily transmissible (12, 13). The mechanisms underlying the development of highly transmissible XDR strains are not fully elucidated. One such mechanism is the induction of efflux pumps, which may lead to high level resistance in mycobacteria (14), without any metabolic compromise. While previous studies described efflux pumps genes and identified mutations in some of these genes (15, 16), efflux pump a transmissible XDR strain have not been described.

Whole genome sequencing using short-reads has elucidated a large number of mutations associated with drug resistance, as well as compensatory mutations, but has limited capacity to resolve large structural variations, gene duplications or variations in repetitive regions (10, 17, 18). Long-read sequencing could provide a more comprehensive understanding of the evolutionary mechanisms underlying the emergence of highly transmissible drug resistant strains (17). In principle, Oxford nanopore MinION sequencing technology offers read lengths that are only limited by the length of DNA presented and produces data in real-time (19). The small size, ease of use and cheap unit cost of the Oxford MinION® nanopore sequencer facilitates successful deployment in resource-limited settings, as has been achieved during the Ebola outbreak in West Africa (20). Although the potential of Oxford MinION® to detect drug resistance mutations in *M. tuberculosis* has been demonstrated, (21) its application for complete *M. tuberculosis* genome assembly has not been reported.

Papua New Guinea (PNG) has a high rate of drug resistant TB in its Western Province (22, 23). We recently characterized a drug resistant tuberculosis outbreak on Daru island, which is driven by a modern Beijing sub-lineage 2.2.1.1 strain (24). Whilst some genetic markers within the strain have been identified (24), the molecular mechanisms responsible for pathogenesis and virulence are not fully elucidated. Genomic regions like proline-glutamate (PE, 99 loci) and proline-proline-glutamate (PPE, 69) genes are routinely excluded in genomic analysis of *M. tuberculosis* due to their repetitive nature and high GC content (9, 24, 25). A recent multinational study utilized 518 sample to study PE/PPE family genes and was able to assemble at least 120 out of the 168 genes for each sample (26).

With signature motifs near the N-terminus of PE/PPE amino acids, these genes are sub classified according to sequence features on the C-terminus. PE genes are divided into PE_PGRS (polymorphic GC-rich sequence, 65 genes) and PE (no distinctive feature, 34 genes) while PPE genes is divided into PPE_MPTR (major polymorphic tandem repeats, 23 genes), PPE_SVP (Gxx-SVPxxW motif, 24 genes), PPE_PPW (PxxPxxW motif, 10 genes) and PPE (no distinctive feature, 12 genes) (27). The existence of these subgroups highlights the diversity in the roles played by these genes (26). PE/PPE gene products are understood to be differentially expressed during infection (28) and have been implicated in immune invasion and virulence (29). We utilized Oxford MinION® to compose a comprehensive draft genome of an XDR strain including variable and repetitive sites like PE/PPE regions. This approach provides insight into the underlying mechanisms of drug resistance and identify key features associated with virulence and transmissibility. The fully assembled genome will serve as an ideal reference for ongoing MDR/XDR outbreak surveillance in Western province, PNG and far north Queensland.

## Materials and methods

### Phenotypic susceptibility testing

The strain was tested for resistance to first and second line drugs including: rifampicin (1·0 μg/ml), isoniazid (0·1 μg/ml; low-level and 0·4 μg/ml; high-level), streptomycin (1·0 μg/ml), ethambutol (5·0 μg/ml), pyrazinamide (100 μg/ml) and second-line drugs amikacin (1·0 μg/ml), capreomycin (2·5 μg/ml), kanamycin (2·5 μg/ml), ethionamide (5·0 μg/ml), ofloxacin (2·0 μg/ml), *p*-aminosalicylic acid (4·0 μg/ml) and cycloserine (50 μg/ml). The automated Bactec Mycobacterial Growth Indicator Tube (MGIT) 960 system (Becton Dickinson, New Jersey, USA) was used for first line drugs and the. Minimal inhibitory concentration assay for second-line drugs was performed using Sensititre® (TREK diagnostic system, Ohio, USA) system for second line drugs.

### DNA extraction and quantification

DNA was extracted from Lowenstien-Jensen (LJ) slopes cultured for 4 weeks at 37°C. DNA isolation was performed using mechanical and chemical methods. Briefly, 5 glass beads (0.7mm, SIGMA) were added to 200μl of PrepMan Ultra sample preparation reagent (Thermo Fisher scientific, Massachusetts, USA). Full loops of culture were added to the reagent and mixed well. The solution was dry heat incubated for 10 minutes at 95°C, followed by bead beating for 40s at 6.0m/s using mini-beadbeater-16 (BioSpec products, Bartlesville, USA). It was then centrifuged for 10 minutes at 13,000rpm before transferring 40μl of the supernatant into another vial. We added 45μl of 3M sodium acetate and 1ml of ice-cold ethanol (96%), centrifuged the solution at maximum speed for 15 minutes and removed the supernatant. We then added 1 ml of 70% ethanol, left it at room temperature for 1 minute, and again removed all the supernatant. The remaining pellet was dried for 15 minutes and re-suspended with 40μl of nuclease free water. The DNA was quantified using Nanodrop 8000 (Thermo fisher scientific, Massachusetts, USA) and Qubit dsDNA HS assay kit (Thermo fisher scientific, Massachusetts, USA)

### MinION® library preparation and sequencing

DNA was purified using 0.4X Agencourt® AMPure® XP beads (Beckman Coulter) and fragment distribution size assessed using Agilent 4200TapeStation (Agilent, UK). Preparation for 1D gDNA library was performed using the SQK-LSK108 manufactures’ instructions. We performed dA-tailing and end-repair using NEBNext Ultra II End-repair/dA-tail module with two step incubation times; 20 minutes each. Then, purification step using 0.7X Agencourt® AMPure® XP beads (Beckman Coulter) was performed according to manufacturers’ instructions. Ligation step was performed using NEB Blunt/ligase master mix module according to manufacturers’ instructions and reaction incubated at room temperature for 20 minutes. Adaptor-ligated DNA was purified using 0.4X Agencourt® AMPure® XP beads (Beckman Coulter) following manufacturers’ instructions but using Oxford Nanopore supplied buffers (adaptor bead binding and elution buffers). The library was ready for MinION® sequencing.

With the MinION MK1B device connected to the computer via a USB3, MinKNOW software (v.1.4.3) was started to perform quality control checks on pore activity and equilibrate the flow cell (FLO-MIN106, version R9.4). The library was combined with reagents supplied by Oxford Nanopore and loaded onto the flow cell following manufacturers’ instructions, choosing a 48h sequencing procedure. Illumina data for the strain was available from our previous study.

### MinION® and Illumina data analysis

Raw files generated by MinKNOW were base called using Albacore (v2.0) to return Oxford Nanopore Technologies (ONT) fastq files. *De novo* genome assembly was performed using Canu (30) and the assembly was improved using consensus with nanopolish (metlylation aware option) (31) and PILON (32). The assembly was circularized using Circulator v1.5.1 (33) and compared with the reference genome H37Rv (NC_000962.3) using MUMMER (34). Genome annotation was performed using the NCBI pipeline (35) and circular representation of the genome viewed using Circos (36). Raw ONT and Illumina reads were mapped to H37Rv using BWA-MEM (37) and assessed genome and base coverage, including PE/PPE families using GATK (DepthOfCoverage) (38). We assessed single nucleotide polymorphisms (SNPs) within PE/PPE genes from nanopore and Illumina reads. In addition, representative raw Illumina reads from four *M. tuberculosis* lineages; Indo-Oceanic, East-Asian (including Beijing lineage), East-African-Indian and Euro-American lineage (including H37Rv) from previous studies (24, 25, 39, 40) were mapped to the draft genome and assessed for large deletions that differentiate the draft genome from other lineages.

### MinION variant and error analysis

Using the reference genome H37Rv (NC_000962.3), single nucleotide polymorphisms (SNPs) and small indels (<5bp) were called from ONT reads using nanopolish (31) and annotated using SnpEff (41). Polymorphisms in known drug resistance genes (including compensatory mutations) were analyzed. MycoBrowser (42) was utilized to analyze mutations in genes that are putatively involved in virulence, efflux pumps and cell transport. Illumina reads of the same strain were mapped onto the reference genome and variants called using GATK (38). Consensus SNPs from the two methods were assessed and ‘non-consensus SNPs’ from ONT reads were considered sequencing errors provided Illumina coverage was greater than 30X. This analysis excluded SNPs in variable regions like PE/PPE genes. Different software to generate variants from ONT and Illumina raw files was combined into an analysis pipeline (Fig. S1).

## Results

In total, 373952 ONT reads passed base calling with N50 read length of 5073bp. The longest read was 33509bp. Genome assembly resulted in one contig of 4404947bp (G+C content 65.5%), with base consistency similar to H37Rv (NC_000962.3) (Fig. S2). NCBI annotation of the genome yielded a total of 3,670 coding DNA sequences (CDS), 45 tRNAs, 3 rRNAs (5S, 16S, 23S) and 3 non-coding RNA (Fig. 1). Mapping of the ONT reads to H37Rv resulted in a coverage of 98.9% at average read depth of 273x (Table S1). Nearly all PE/PPE genes (167/168) were completely assembled with 100% coverage; only one (*wag22;* RV1759c) had incomplete (88%) coverage in our assembly relative to H37Rv. The average ONT read depth of PE/PPE genes was 299.87 (IQR 285.91-311.36) (Fig. S3). Only 54.3% (92/168) of the PE/PPE genes were completely assembled from Illumina contigs at average sequence depth of x46.3 (Fig. S4). No Illumina reads covered the PE_PGRS sub-family genes *wag22* and PE_PGRS57.

**Fig. 1:**
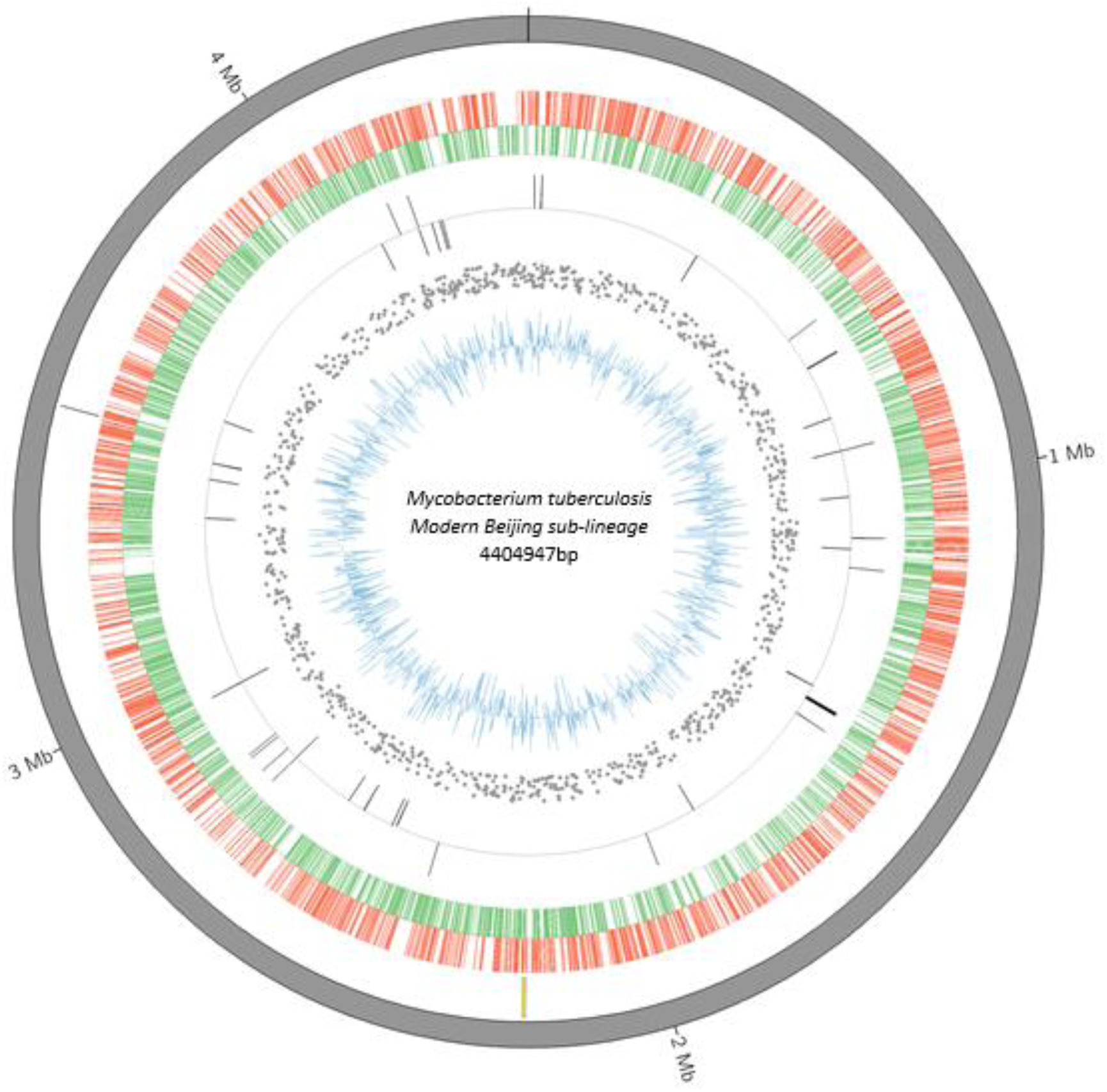
Circular representation of the drafted XDR genome from Western Province, Papua New Guinea (modern Beijing sub-lineage 2.2.1.1 strain) with gene annotations. From the inner to outer ring: Blue - GC content of coding sequence; Scattered grey - SNPs relative to H37Rv; Grey bars - reverse and forward *rRNA*; Green - reverse coding sequences; Red - forward coding sequences; Yellow bars - unique regions compared to H37Rv; Grey outside ring - assembled contig (grey). Mb-million base pairs; XDR – extensively drug resistant

We first evaluated SNP calling in the non-repetitive part of the reference genome by excluding variable (PE/PPE) regions. A total of 1254 SNPs and 122 small indels were called from ONT reads while 1098 SNPs and 105 small indels were called from Illumina reads when mapped to H37Rv. Of these, 1095 SNPs (574 non-synonymous and 402 synonymous) and 87 small indels were identified by both approaches. 118 of 159 SNPs and 23 of 35 indels identified from MinION® but not Illumina were in regions of low Illumina coverage (<30x coverage). The remaining 41 and 12 SNPs and Indels respectively are potentially due to systematic base-calling errors (3.2% and 9.8 %). Three (0.2%) SNPs and 18 (14.7%) indels were identified by Illumina, but not by MinION® sequencing. As a conservative estimate considering all 194 ONT only calls (159 SNPs and 35 indels) as false positives and 21 Illumina only calls (3 SNPs and 18 Indels) as false negative, the error rate after consensus calling from ONT was 0.0048%. If we ignore inconsistent calls where Illumina coverage was low then the estimated error rate for base calling from ONT was reduced to 0.0036%

Given the likely importance of the variable PE/PPE genes in strain evolution we assessed the ability of MinION® and Illumina to call SNPs in this class of genes separately. From nanopore assembly, 158 SNPs were identified from 70 PE/PPE genes (42 PE and 28 PPE) with 88 SNPs (55.6%) identified from the PE_PGRS sub-family (Table S2). From the Illumina assembly, 124 SNPs from 45 PE/PPE genes (25 PE and 20 PPE) were identified of which 31 SNPs (25%) were from PE_PGRS sub-family. There were 81 SNPs (from 42 PE/PPE genes) overlapping between ONT and Illumina with PPE54 having the highest number of overlapped SNPs (9) identified within one gene.

Phenotypic drug susceptibility results revealed the isolate to be extensively drug resistant with susceptibility to only amikacin, kanamycin, para-aminosalicyclic acid (PAS) and cycloserine. Table 1 reflects phenotypic resistance, as well as mutations in genes known to confer drug resistance to first and second line drugs and recognized compensatory mutations. Genotypic drug resistance profiles concurred with phenotypic results. While 10 mutations were identified in seven genes that encode trans-membrane efflux pumps and transporter proteins (Table 2). Table 3 shows 16 SNPs identified in genes that encode virulence proteins; 8 (50%) were from the *mce*-gene family and a mutation within *mycP1* (p.Thr238Ala) was also noted. In addition, 27 SNPs were identified in three genes families involved in cell wall synthesis, with 17 in *fadD*, 4 in *pks* and 3 in *mmp* gene families (Table S3).

**Table 1:** Mutations in candidate drug resistance genes identified from ONT assembly of XDR Beijing sub-lineage 2.2.1.1 strain ONT – Oxford Nanopore Technologies; XDR – Extensively drug resistant; PZA – Pyrazinamide; PAS - Para-amino salicylic acid

| Drug | Invitro phenotype | Investigated genes | Genes (mutation) |
| --- | --- | --- | --- |
| Isoniazid | Resistant | *fabG1-inhA, inhA, katG, ndh, furA, oxyR, aphC, fadE24, srmR, kasA, mshA* | *fabG1-inhA* (C-15T) |
|  |  |  | *inhA* (p.Ile21Val) |
|  |  |  | *ndh* (delG304) |
| Rifampicin | Resistant | *rpoB, rpoC, rpoA, rpoD* | *rpoB* (p.Ser450Leu) |
|  |  |  | *rpoC* (p.Val483Gly) |
| Ethambutol | Resistant | *embB, embC, embA, ubiA, embR, iniA, iniC, manB* | *embB* (p.Met306Val) |
| Pyrazinamide | Resistant | *pncA, rpsA, panD* | *pncA*(p.Tyr103Asp) |
| Streptomycin | Resistant | *rpsL, gidB, rrs* | *rpsL* (p.Lys43Arg) |
|  |  |  | *gidB* (p.Leu91Pro) |
| Ethionamide | Resistant | *fabG1-inhA, ethA, ethR* | *fabG1-inhA*(C-15T) |
| Fluoroquinolone | Resistant | *gyrA, gyrB* | *gyrA* (p.Asp94Gly) |
| Amikacin | Susceptible | *rrs, whib7, gidB* | nil |
| Kanamycin | Susceptible | *Rv2417c-eis, whiB7, rrs, gidB* | nil |
| Capreomycin | Resistant | *tlyA, whiB7, rrs, gidB* | *tlyA* (ins397C) |
| PAS | Susceptible | *ribD, thyA, dfrA, folC* | nil |
| Cycloserine | Susceptible | *alr, ddl, cycA* | nil |

**Table 2.**
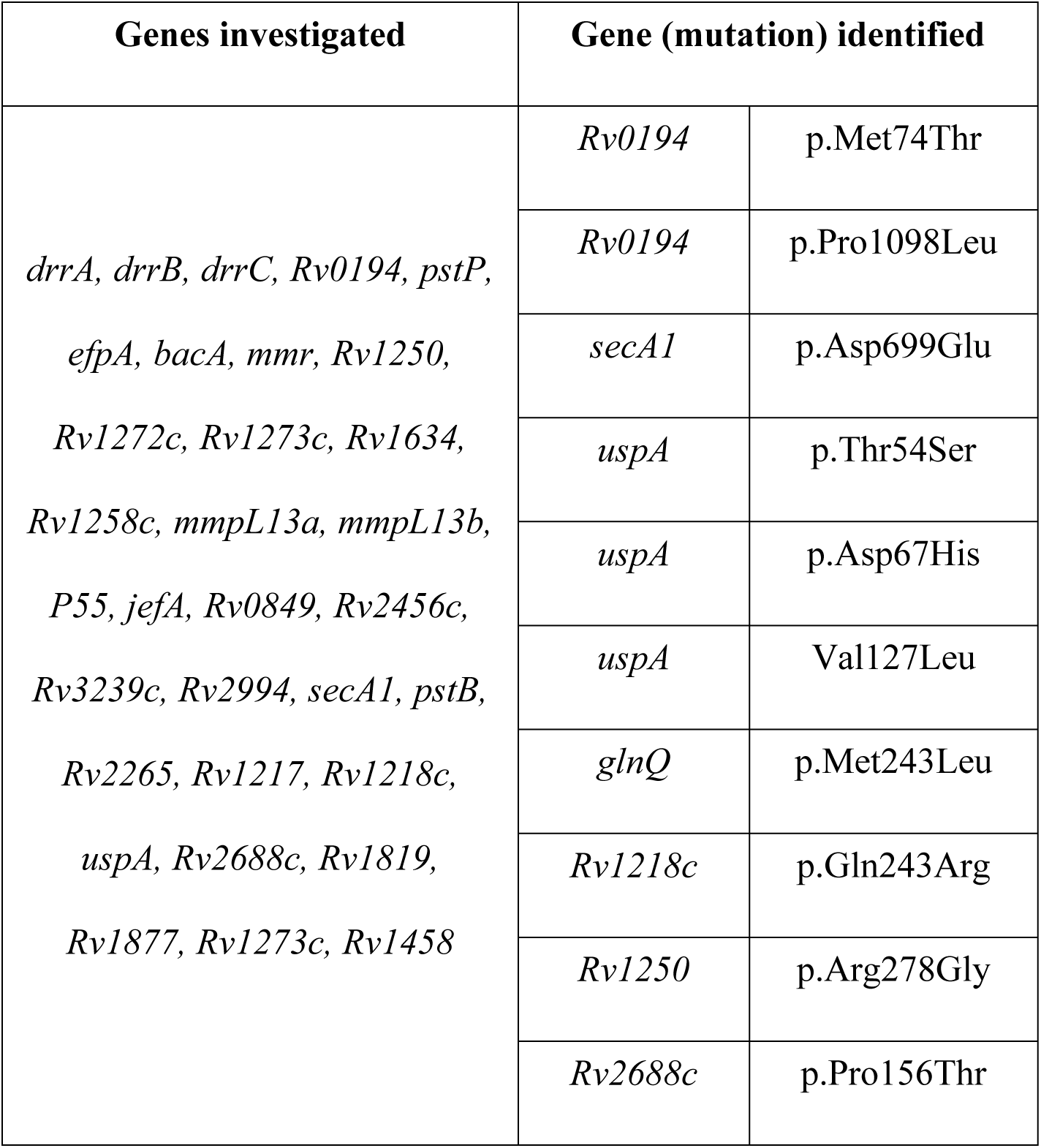
Mutations in putative efflux pump/transporter genes identified from ONT assembly of XDR Beijing sub-lineage 2.2.1.1 strain ONT – Oxford Nanopore Technologies; XDR - Extensively drug resistant

**Table 3:** Mutations identified in genes that encode for potential virulence proteins in the XDR Beijing sub-lineage 2.2.1.1 strain XDR - Extensively drug resistant

| Gene | Nucleotide change | Position | Amino acid |
| --- | --- | --- | --- |
| *mak* | T->C | 154283 | p.Ser18Pro |
| *mce1A* | T->G | 199470 | p.Ser313Ala |
| *mce1D* | T->C | 203038 | p.Ile188Thr |
| *mce1F* | T->C | 206339 | p.Leu370Pro |
| *mce2A* | T->C | 686972 | p.Phe51Ser |
| *mce2F* | A->G | 694531 | p.Asn432Ser |
| *Rv0634c* | A->G | 731015 | p.Tyr7His |
| *mazF3* | G->A | 1230778 | p.Thr65Ile |
| *ephB* | G->T | 2191498 | p.Gly158Trp |
| *mce3A* | G->A | 2209465 | p.Ala47Thr |
| *mce3F* | C->A | 2216443 | p.Ala396Glu |
| *cstA* | C->A | 3428917 | p.Arg559Ser |
| *mce4C* | G->T | 3916386 | pArg191Ser |
| *proV* | T->C | 4204168 | p.Asn84Asp |
| *proX* | A->G | 4205120 | p.Leu85Pro |
| *mycP1* | T->C | 4364046 | Thr238Ala |

Mapping of raw sequence reads from representative lineage 1 (Indo-Oceanic), 2 (East Asian, including ancient and modern Beijing), 3 (East-African-Indian) and 4 (Euro-American, including H37Rv) strains identified a 4490bp (2207042-2211532) region absent in Euro-American lineage strains (Fig. 2). Three previously assembled Beijing genomes from PacBio long reads identified the same region (40) (Fig. S5). Like is previous studies (43-45), annotation of this region was revealed to span 7 complete genes that encode proteins that include a nicotinamide adenine dinucleotide phosphate (NADP)-dependent oxidoreductase, an iron-regulated elongation factor (Tu), a PE-family protein while four genes encode uncharacterized proteins (Fig. 2).

**Fig. 2:**
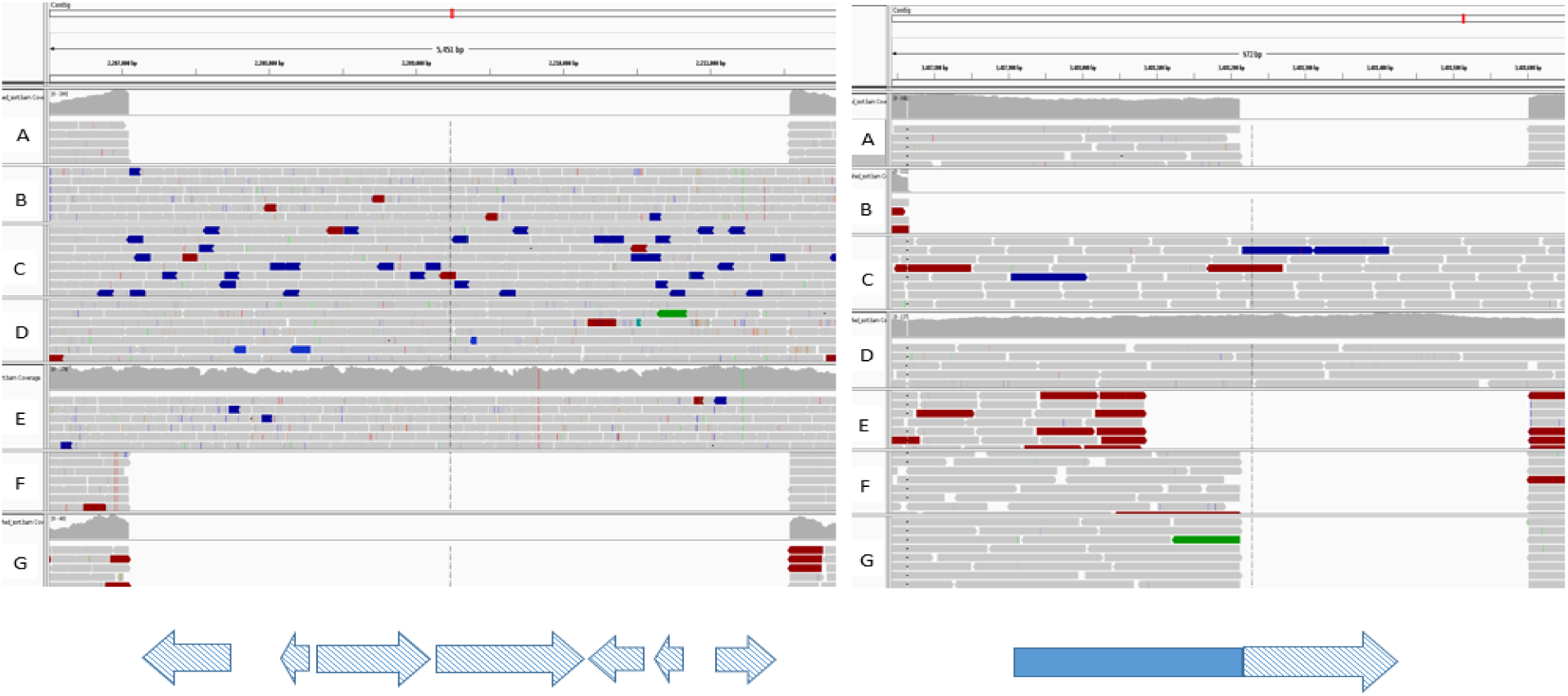
Integrative Genomic Viewer (IGV) of Illumina reads from different *M. tuberculosis* lineages (A-*M. tuberculosis* H37Rv, B-Indo-oceanic, C-Ancient Beijing, D-Modern Beijing, E-East African Indian, F&G-Euro American) mapped on the ONT draft genome highlighting the large insertions within the draft genome. On the left-4490bp insertion (2207042-2211532) spanning 7 annotated genes (blue checked arrows-NADH-dependent oxidoreducatse, Iron-regulated elongation factor-tu, PE family protein and hypothetical/uncharacterized proteins). On the right-390bp insertion relative to H37Rv (3488211-3488601) spanning 323bp (checked) at the end of a 654bp coding sequence. ONT – Oxford Nanopore Technologies

A second smaller insertion (390bp, 3488211-3488601) was identified among Beijing lineage genomes relative to H37Rv and Euro-American lineages but varied in size as it was 835bp with respect to Indo-oceanic and East-African-Indian lineages (Fig. 2). Annotation showed a 654bp gene (3487881-3488534) in this region had part of the insertion sequence, 323bp (3488211-3488534) towards the end. Phyre2 protein modelling (46) of the gene sequence with the insertion revealed PE8-PPE15 as template to construct to predict the protein structure as a PPE family protein (79% sequence modelled, 100% confidence) consisting of 73% alpha helices (Fig. S6). A blast search of this gene sequence revealed a 50% query coverage to four *Mycobacterium tuberculosis* H37Rv genomes (100% identification) and 100% query coverage to 55 *Mycobacterium tuberculosis*, Lineage 2 genomes (Table S4).

## Discussion

In this study, we utilized Oxford Nanopore MinION sequencing to assemble a comprehensive genome of a strain that is responsible for a drug resistant outbreaks in the Western Province of Papua New Guinea (24). The complete circular genome of this modern Beijing sub-lineage 2.2.1.1 strain revealed genetic determinants of drug resistance against first and second line TB drugs. Nanopore sequencing allowed us to assemble highly variable PE/PPE gene families with great fidelity. PE/PPE genes are thought to encode surface-associated cell wall proteins that may provide antigenic diversity and affect host immunity (28). Nanopore technology has been previously used to improve genome assemblies and resolution of repeat-rich regions in S*almonella typhi* and *Escerischia coli* (18, 31) but not yet with *Mycobacterium tuberculosis*.

Twice as many SNPs in PE_PGRS genes were identified as by ONT Minion versus Illumina sequencing (88 vs 31). PE_PGRS gene mutations were under-represented in short-read sequencing, possibly due to their extra GC containing motifs that impact on the sequencing. Previous studies have identified a higher number of mutations within the PE_PGRS sub-family compared to other PE subfamilies, and attribute it to their involvement in antigenic variation and immune evasion from exposure to host immune system (26, 47). The *wag22* gene was also better represented by nanopore sequencing, with 88% coverage compared to no coverage from Illumina reads. Besides the high GC content, the difficulty of sequencing *wag22* has been further attributed to deletions at the beginning of the open reading frame (47). Unsurprisingly, PPE54 which is a member of PPE_MPTR (major polymorphic tandem repeat) sub-family had the highest number of mutations from ONT sequence. Previous studies have shown it to be involved in the ‘arrest’ of phagosome maturation to allow survival of the bacteria in the macrophages due to its long amino acid length at the C-terminal (48, 49). It has been postulated that PPE54 gene mutations may also play a role in development of isoniazid, rifampicin and ethambutol resistance (50), but we were unable to verify this given the presence of well-characterized drug resistance mutations.

There is growing interest in using Oxford Nanopore Technologies devices for real-time clinical utility as a cheap point-of-care TB diagnostic, with accurate identification of antimicrobial resistance profiles. Sequencing for drug resistance mutations directly from clinical samples has been completed within a 24 hours in patients with sputum smear-positive tuberculosis (21). In our study, nanopore sequencing of an XDR strain fully identified its drug resistance profile with complete phenotypic concordance. We were able to identify all relevant first line and second line drug resistance conferring SNPs using Oxford nanopore MinION seqeuncing. Compensatory mutations also detected in genes like *rpoC* (p.Val483Gly) and *ndh* (c.304delG) are thought to ameliorate the fitness cost associated with the XDR phenotype (51, 52).

We also identified mutations in efflux pumps and transporter proteins, which might contribute to resistance phenotypes (53). Mutations in transporter proteins like ABC (ATP binding cassette) and MFS (Major Facilitator Super family) have been associated with drug resistance (54, 55). For example, mutations leading to overexpression of the ABC transporter Rv0194 leads to increased export of multiple drugs like streptomycin, vancomycin, and tetracycline (56). We identified two mutations in *uspA*, which is part of the three gene operon *uspABC* that encodes membrane-spanning subunits transporting amino-sugar substrates across the cell wall (57). Mutations found in Rv0194 have been associated with resistance to beta lactams antibiotics (56) and it’s over expression with an XDR phenotype (58). Further research needs to be done to explore the association between efflux pump mutations, pump activity and drug resistance.

Mutations in co-localized genes like; *mmpL*, *pks* and *fadD* have been considered to a play a compensatory role to restore the fitness of drug resistant strains (59, 60) especially for drugs that target biosynthesis pathways of the mycobacterial cell wall like isoniazid, ethionamide (5) and ethambutol (61). *M. tuberculosis* contains 13 genes that encode *mmpL* proteins and 16 genes that encode polyketide synthases (*pks*) proteins that are involved in lipopolysaccharide and complex lipid biosynthesis. The functional cross talk between *pks* and *fadD* genes has been demonstrated in studies that showed how *pks13* and *fad32* form specific substrates that are precursors of mycolic acid biosynthesis (62). We identified mutations in all three genes although there was greater mutation variability within *fadD* genes. It remains to be determined how mutations in these co-localized genes influence cell wall lipid biosynthesis.

Insight into the factors that influence mycobacterial virulence is important for better appreciation of microbial pathogenesis and the identification of new treatment options. SNPs in mammalian cell entry (*mce*) genes were prominent. Mce-family proteins are proposed to be involved in invasion and persistence of *M. tuberculosis* in host macrophages (63). This is related to the ability of these cell surface proteins to mediate bacterial uptake by mammalian cells, similar to those stimulated by invasive enteric bacteria (64). It has been demonstrated that a mutant *mce1A M. tuberculosis* strain is “hyper virulent” in mice (65). Comparative analysis among different *M. tuberculosis* strains could unveil characteristics related to host adaption (66).

Although *M. tuberculosis* has relatively limited genetic variation compared to other pathogenic bacteria, there is strain-related phenotypic variation in the protection provided by Bacille Calmette-Guerin (BCG) vaccination and clinical outcome (67). We didn’t find any large indel (>1kb) unique to the study strain but identified a large region (4490bp) with 7 coding sequences present in the draft genome and other reference genomes (lineage 1, 2 and 3) but absent in lineage 4 genomes including H37Rv. These have been previously described (43-45) and this further demonstrates the limitations of using H37Rv as the universal reference strain. The second smaller region within the draft genome but with variable sizes among the reference genomes highlights evidence of independent structural rearrangement among the different lineages. The identified PPE family protein unique to Beijing lineage 2 could contribute to phenotypic characteristics of this lineage. Such a comparative approach provides an opportunity to study lineage and strain specific differences, especially in the advent of long read sequencing with enhanced resolution of variable parts of the genome.

In conclusion, the assembly of a complete genome of a XDR “epidemic strain” using nanopore technology did not only provide proof of principle for future deployment of this technology in settings endemic for drug resistant tuberculosis but it also demonstrated the use of this technology in further understanding of *M. tuberculosis* genetics. It characterized the drug resistance profile and potential virulence factors found in this strain, and provided a reference strain for future genome assembly and mapping.

## Data bibliography

1. NCBI project accession number PRJNA386696 (2018)
2. GenBank accession numbers AP018034 (HN-205), AP018035 (HN-321), and AP018036(HN-506)-2017

## Ethic statement

The isolate was selected from a previous study (24) with ethics clearance from the University of Queensland and the Papua New Guinea Medical Research Advisory Committee to perform detailed whole genome sequencing.

## Acknowledgements

We thank the Papua New Guinea National Tuberculosis Program, Provincial Health Staff (Western Province) and staff of Queensland Mycobacterium Reference Laboratory for their assistance. This research was supported by the use of the NeCTAR Research Cloud, by QCIF and by the University of Queensland’s Research Computing Centre (RCC). The NeCTAR Research Cloud is a collaborative Australian research platform supported by the National Collaborative Research Infrastructure Strategy.

## Supplementary

**Fig. S1**: A scheme of workflow to generate variants from Oxford Nanopore technologies (ONT) and Illumina reads

**Fig. S2**: Dot plot of sequence accuracy between the draft Beijing sub-lineage 2.2.1.1 strain genome and the reference genome H37Rv. The red dots show a forward orientation while blue dots show reverse orientation

**Fig. S3**: Plot of sequence and depth coverage for the assembly of 168 PE/PPE family genes using Oxford minion® reads. The marks on the x-axis represent the different 168 PE/PPE family genes while y-axis respective their percentage assembly coverage (red), and read depth (blue). Sample was sequenced to an average of 273x depth

**Fig. S4**: Plot of sequence and depth coverage for the assembly of 168 PE/PPE family genes using Illumina reads. The marks on the x-axis represent the different 168 PE/PPE family genes while y-axis respective their percentage assembly coverage (red), and read depth (blue). Sample was sequenced to average 46.3x depth

**Fig. S5**: Dot plot of ‘unique region’ accuracy between the draft Beijing sub-lineage 2.2.1.1 strain (x-axis) genome and one of the Beijing reference genomes assembled from PacBio reads (y-axis), genebank accession number AP018035 (HN-321).

**Fig. S6**: Protein structure of the PPE family protein predicted using Phyre2 for a 654bp gene sequence with an end 323bp insertion identified within the assembled genome but absent in the reference genome H37Rv.

**Table S1**: Details of Oxford Nanopore Technologies (ONT) reads utilized and assembled genome

**Table S2**: Number of SNPs and average base coverage identified from different assembled PE/PPE family genes from ONT and Illumina reads

**Table S3**: Mutations in genes involved in cell wall biosynthesis identified in the Beijing sub-lineage 2.2.1.1 strain

**Table S4**: Blast search results of the gene sequence with insertion confirming the uniqueness of insertion sequence among lineage 2 genomes

## References

1. World Health Organization. GLOBAL TUBERCULOSIS REPORT 2016. Switzerland: World Health Organization, Geneva, 2017.

2. Abubakar I, Zignol M, Falzon D, Raviglione M, Ditiu L, Masham S, et al. Drug-resistant tuberculosis: time for visionary political leadership. The Lancet Infectious diseases. 2013;13(6):529–39.

3. Dheda K, Gumbo T, Maartens G, Dooley KE, McNerney R, Murray M, et al. The epidemiology, pathogenesis, transmission, diagnosis, and management of multidrug-resistant, extensively drug-resistant, and incurable tuberculosis. The Lancet Respiratory medicine. 2017.

4. Udwadia ZF. MDR, XDR, TDR tuberculosis: ominous progression. Thorax. 2012;67(4):286–8.

5. Zhang Y, Yew WW. Mechanisms of drug resistance in Mycobacterium tuberculosis: update 2015. The international journal of tuberculosis and lung disease: the official journal of the International Union against Tuberculosis and Lung Disease. 2015;19(11):1276–89.

6. Eldholm V, Monteserin J, Rieux A, Lopez B, Sobkowiak B, Ritacco V, et al. Four decades of transmission of a multidrug-resistant Mycobacterium tuberculosis outbreak strain. Nature communications. 2015;6:7119.

7. Ioerger TR, Feng Y, Chen X, Dobos KM, Victor TC, Streicher EM, et al. The non-clonality of drug resistance in Beijing-genotype isolates of Mycobacterium tuberculosis from the Western Cape of South Africa. BMC genomics. 2010;11:670.

8. Didelot X, Walker AS, Peto TE, Crook DW, Wilson DJ. Within-host evolution of bacterial pathogens. Nature reviews Microbiology. 2016;14(3):150–62.

9. Cohen KA, Abeel T, Manson McGuire A, Desjardins CA, Munsamy V, Shea TP, et al. Evolution of Extensively Drug-Resistant Tuberculosis over Four Decades: Whole Genome Sequencing and Dating Analysis of Mycobacterium tuberculosis Isolates from KwaZulu-Natal. PLoS medicine. 2015;12(9):e1001880.

10. Casali N, Nikolayevskyy V, Balabanova Y, Harris SR, Ignatyeva O, Kontsevaya I, et al. Evolution and transmission of drug-resistant tuberculosis in a Russian population. Nature genetics. 2014;46(3):279–86.

11. Casali N, Nikolayevskyy V, Balabanova Y, Ignatyeva O, Kontsevaya I, Harris SR, et al. Microevolution of extensively drug-resistant tuberculosis in Russia. Genome research. 2012;22(4):735–45.

12. McBryde ES, Meehan MT, Doan TN, Ragonnet R, Marais BJ, Guernier V, et al. The risk of global epidemic replacement with drug-resistant Mycobacterium tuberculosis strains. International journal of infectious diseases: IJID: official publication of the International Society for Infectious Diseases. 2017;56:14–20.

13. Marais BJ, Mlambo CK, Rastogi N, Zozio T, Duse AG, Victor TC, et al. Epidemic spread of multidrug-resistant tuberculosis in Johannesburg, South Africa. Journal of clinical microbiology. 2013;51(6):1818–25.

14. Schmalstieg AM, Srivastava S, Belkaya S, Deshpande D, Meek C, Leff R, et al. The antibiotic resistance arrow of time: efflux pump induction is a general first step in the evolution of mycobacterial drug resistance. Antimicrobial agents and chemotherapy. 2012;56(9):4806–15.

15. Garima K, Pathak R, Tandon R, Rathor N, Sinha R, Bose M, et al. Differential expression of efflux pump genes of Mycobacterium tuberculosis in response to varied subinhibitory concentrations of antituberculosis agents. Tuberculosis. 2015;95(2):155–61.

16. Ali A, Hasan Z, McNerney R, Mallard K, Hill-Cawthorne G, Coll F, et al. Whole genome sequencing based characterization of extensively drug-resistant Mycobacterium tuberculosis isolates from Pakistan. PloS one. 2015;10(2):e0117771.

17. Didelot X, Bowden R, Wilson DJ, Peto TEA, Crook DW. Transforming clinical microbiology with bacterial genome sequencing. Nature reviews Genetics. 2012;13(9):601–12.

18. Ashton PM, Nair S, Dallman T, Rubino S, Rabsch W, Mwaigwisya S, et al. MinION nanopore sequencing identifies the position and structure of a bacterial antibiotic resistance island. Nature biotechnology. 2015;33(3):296–300.

19. Laver T, Harrison J, O’Neill PA, Moore K, Farbos A, Paszkiewicz K, et al. Assessing the performance of the Oxford Nanopore Technologies MinION. Biomolecular detection and quantification. 2015;3:1–8.

20. Quick J, Loman NJ, Duraffour S, Simpson JT, Severi E, Cowley L, et al. Real-time, portable genome sequencing for Ebola surveillance. Nature. 2016;530(7589):228–32.

21. Votintseva AA, Bradley P, Pankhurst L, Del Ojo Elias C, Loose M, Nilgiriwala K, et al. Same-Day Diagnostic and Surveillance Data for Tuberculosis via Whole-Genome Sequencing of Direct Respiratory Samples. Journal of clinical microbiology. 2017;55(5):1285–98.

22. Organization WH. Global tuberculosis control: WHO report. Geneva: Global Tuberculosis Programme. p. 15 volumes.

23. Aia P, Kal M, Lavu E, John LN, Johnson K, Coulter C, et al. The Burden of Drug-Resistant Tuberculosis in Papua New Guinea: Results of a Large Population-Based Survey. PloS one. 2016;11(3):e0149806.

24. Bainomugisa A, Lavu E, Hiashiri S, Majumdar S, Honjepari A, Moke R, et al. Multi-clonal evolution of multi-drug-resistant/extensively drug-resistant Mycobacterium tuberculosis in a high-prevalence setting of Papua New Guinea for over three decades. Microbial genomics. 2018.

25. Merker M, Blin C, Mona S, Duforet-Frebourg N, Lecher S, Willery E, et al. Evolutionary history and global spread of the Mycobacterium tuberculosis Beijing lineage. Nature genetics. 2015;47(3):242–9.

26. Phelan JE, Coll F, Bergval I, Anthony RM, Warren R, Sampson SL, et al. Recombination in pe/ppe genes contributes to genetic variation in Mycobacterium tuberculosis lineages. BMC genomics. 2016;17:151.

27. Fishbein S, van Wyk N, Warren RM, Sampson SL. Phylogeny to function: PE/PPE protein evolution and impact on Mycobacterium tuberculosis pathogenicity. Molecular microbiology. 2015;96(5):901–16.

28. Mohareer K, Tundup S, Hasnain SE. Transcriptional regulation of Mycobacterium tuberculosis PE/PPE genes: a molecular switch to virulence? Journal of molecular microbiology and biotechnology. 2011;21(3-4):97–109.

29. Singh KK, Zhang X, Patibandla AS, Chien P, Jr., Laal S. Antigens of Mycobacterium tuberculosis expressed during preclinical tuberculosis: serological immunodominance of proteins with repetitive amino acid sequences. Infection and immunity. 2001;69(6):4185–91.

30. Koren S, Walenz BP, Berlin K, Miller JR, Bergman NH, Phillippy AM. Canu: scalable and accurate long-read assembly via adaptive k-mer weighting and repeat separation. Genome research. 2017;27(5):722–36.

31. Loman NJ, Quick J, Simpson JT. A complete bacterial genome assembled de novo using only nanopore sequencing data. Nature methods. 2015;12(8):733–5.

32. Walker BJ, Abeel T, Shea T, Priest M, Abouelliel A, Sakthikumar S, et al. Pilon: an integrated tool for comprehensive microbial variant detection and genome assembly improvement. PloS one. 2014;9(11):e112963.

33. Hunt M, Silva ND, Otto TD, Parkhill J, Keane JA, Harris SR. Circlator: automated circularization of genome assemblies using long sequencing reads. Genome biology. 2015;16:294.

34. Kurtz S, Phillippy A, Delcher AL, Smoot M, Shumway M, Antonescu C, et al. Versatile and open software for comparing large genomes. Genome biology. 2004;5(2):R12.

35. Angiuoli SV, Gussman A, Klimke W, Cochrane G, Field D, Garrity G, et al. Toward an online repository of Standard Operating Procedures (SOPs) for (meta)genomic annotation. Omics: a journal of integrative biology. 2008;12(2):137–41.

36. Krzywinski M, Schein J, Birol I, Connors J, Gascoyne R, Horsman D, et al. Circos: an information aesthetic for comparative genomics. Genome research. 2009;19(9):1639–45.

37. Li H. Aligning sequence reads, clone sequences and assembly contigs with BWA-MEM. arXiv. 2013;:1303.3997v1 [q-bio.GN].

38. McKenna A, Hanna M, Banks E, Sivachenko A, Cibulskis K, Kernytsky A, et al. The Genome Analysis Toolkit: a MapReduce framework for analyzing next-generation DNA sequencing data. Genome research. 2010;20(9):1297–303.

39. Guerra-Assuncao JA, Crampin AC, Houben RM, Mzembe T, Mallard K, Coll F, et al. Large-scale whole genome sequencing of M. tuberculosis provides insights into transmission in a high prevalence area. eLife. 2015;4.

40. Wada T, Hijikata M, Maeda S, Hang NTL, Thuong PH, Hoang NP, et al. Complete Genome Sequences of Three Representative Mycobacterium tuberculosis Beijing Family Strains Belonging to Distinct Genotype Clusters in Hanoi, Vietnam, during 2007 to 2009. Genome announcements. 2017;5(27).

41. Cingolani P, Platts A, Wang le L, Coon M, Nguyen T, Wang L, et al. A program for annotating and predicting the effects of single nucleotide polymorphisms, SnpEff: SNPs in the genome of Drosophila melanogaster strain w1118; iso-2; iso-3. Fly. 2012;6(2):80–92.

42. Kapopoulou A, Lew JM, Cole ST. The MycoBrowser portal: a comprehensive and manually annotated resource for mycobacterial genomes. Tuberculosis. 2011;91(1):8–13.

43. Periwal V, Patowary A, Vellarikkal SK, Gupta A, Singh M, Mittal A, et al. Comparative whole-genome analysis of clinical isolates reveals characteristic architecture of Mycobacterium tuberculosis pangenome. PloS one. 2015;10(4):e0122979.

44. O’Toole RF, Gautam SS. Limitations of the Mycobacterium tuberculosis reference genome H37Rv in the detection of virulence-related loci. Genomics. 2017.

45. Gautam SS, Mac Aogain M, Bower JE, Basu I, O’Toole RF. Differential carriage of virulence-associated loci in the New Zealand Rangipo outbreak strain of Mycobacterium tuberculosis. Infectious diseases. 2017;49(9):680–8.

46. Kelley LA, Mezulis S, Yates CM, Wass MN, Sternberg MJ. The Phyre2 web portal for protein modeling, prediction and analysis. Nature protocols. 2015;10(6):845–58.

47. Copin R, Coscolla M, Seiffert SN, Bothamley G, Sutherland J, Mbayo G, et al. Sequence diversity in the pe_pgrs genes of Mycobacterium tuberculosis is independent of human T cell recognition. mBio. 2014;5(1):e00960–13.

48. Sampson SL. Mycobacterial PE/PPE proteins at the host-pathogen interface. Clinical & developmental immunology. 2011;2011:497203.

49. Brodin P, Poquet Y, Levillain F, Peguillet I, Larrouy-Maumus G, Gilleron M, et al. High content phenotypic cell-based visual screen identifies Mycobacterium tuberculosis acyltrehalose-containing glycolipids involved in phagosome remodeling. PLoS pathogens. 2010;6(9):e1001100.

50. Cui ZJ, Yang QY, Zhang HY, Zhu Q, Zhang QY. Bioinformatics Identification of Drug Resistance-Associated Gene Pairs in Mycobacterium tuberculosis. International journal of molecular sciences. 2016;17(9).

51. Comas I, Borrell S, Roetzer A, Rose G, Malla B, Kato-Maeda M, et al. Whole-genome sequencing of rifampicin-resistant Mycobacterium tuberculosis strains identifies compensatory mutations in RNA polymerase genes. Nature genetics. 2012;44(1):106–10.

52. Bainomugisa A, Lavu E, Hiashiri S, Majumdar S, Honjepari A, Moke R, et al. Multi-clonal evolution of MDR/XDR M. tuberculosis in a high prevalence setting in Papua New Guinea over three decades. bioRxiv. 2017.

53. Kanji A, Hasan R, Zhang Y, Shi W, Imtiaz K, Iqbal K, et al. Increased expression of efflux pump genes in extensively drug-resistant isolates of Mycobacterium tuberculosis. International journal of mycobacteriology. 2016;5 Suppl 1:S150.

54. Liu J, Takiff HE, Nikaido H. Active efflux of fluoroquinolones in Mycobacterium smegmatis mediated by LfrA, a multidrug efflux pump. Journal of bacteriology. 1996;178(13):3791–5.

55. De Rossi E, Arrigo P, Bellinzoni M, Silva PA, Martin C, Ainsa JA, et al. The multidrug transporters belonging to major facilitator superfamily in Mycobacterium tuberculosis. Molecular medicine. 2002;8(11):714–24.

56. Danilchanka O, Mailaender C, Niederweis M. Identification of a novel multidrug efflux pump of Mycobacterium tuberculosis. Antimicrobial agents and chemotherapy. 2008;52(7):2503–11.

57. Fullam E, Prokes I, Futterer K, Besra GS. Structural and functional analysis of the solute-binding protein UspC from Mycobacterium tuberculosis that is specific for amino sugars. Open biology. 2016;6(6).

58. Kanji A, Hasan R, Ali A, Zaver A, Zhang Y, Imtiaz K, et al. Single nucleotide polymorphisms in efflux pumps genes in extensively drug resistant Mycobacterium tuberculosis isolates from Pakistan. Tuberculosis. 2017;107(Supplement C):20–30.

59. Zhang H, Li D, Zhao L, Fleming J, Lin N, Wang T, et al. Genome sequencing of 161 Mycobacterium tuberculosis isolates from China identifies genes and intergenic regions associated with drug resistance. Nature genetics. 2013;45(10):1255–60.

60. Kuan CS, Chan CL, Yew SM, Toh YF, Khoo JS, Chong J, et al. Genome Analysis of the First Extensively Drug-Resistant (XDR) Mycobacterium tuberculosis in Malaysia Provides Insights into the Genetic Basis of Its Biology and Drug Resistance. PloS one. 2015;10(6):e0131694.

61. Birch HL, Alderwick LJ, Bhatt A, Rittmann D, Krumbach K, Singh A, et al. Biosynthesis of mycobacterial arabinogalactan: identification of a novel alpha(1–>3) arabinofuranosyltransferase. Molecular microbiology. 2008;69(5):1191–206.

62. Gavalda S, Leger M, van der Rest B, Stella A, Bardou F, Montrozier H, et al. The Pks13/FadD32 crosstalk for the biosynthesis of mycolic acids in Mycobacterium tuberculosis. The Journal of biological chemistry. 2009;284(29):19255–64.

63. Casali N, Riley LW. A phylogenomic analysis of the Actinomycetales mce operons. BMC genomics. 2007;8:60.

64. Bliska JB, Galan JE, Falkow S. Signal transduction in the mammalian cell during bacterial attachment and entry. Cell. 1993;73(5):903–20.

65. Shimono N, Morici L, Casali N, Cantrell S, Sidders B, Ehrt S, et al. Hypervirulent mutant of Mycobacterium tuberculosis resulting from disruption of the mce1 operon. Proceedings of the National Academy of Sciences of the United States of America. 2003;100(26):15918–23.

66. Cubillos-Ruiz A, Morales J, Zambrano MM. Analysis of the genetic variation in Mycobacterium tuberculosis strains by multiple genome alignments. BMC research notes. 2008;1:110.

67. Gagneux S, Small PM. Global phylogeography of Mycobacterium tuberculosis and implications for tuberculosis product development. The Lancet Infectious diseases. 2007;7(5):328–37.

